# Thermal biology of two sympatric Lacertids lizards (*Lacerta diplochondrodes* and *Parvilacerta parva*) from Western Anatolia

**DOI:** 10.1101/2021.03.16.435405

**Authors:** Mehmet Kürşat Şahin, Arda Cem Kuyucu

## Abstract

Sympatric lizard species differing in morphology present convenient models for studying the differentiation in thermal behavior and the role of morphological differences in thermal biology. Here we studied the thermal biology of two sympatric lizard species which occur together sympatrically in western Anatolia, Frig Valley. These two species differ in body size, with the larger *Lacerta diplochondrodes* and smaller *Parvilacerta parva*. Field body temperatures of the individuals belonging to both species were recorded in the activity period. Additionally, several environmental parameters including solar radiation, substrate temperature, air temperature and wind speed were also monitored to investigate the relative effect of these abiotic parameters on thermal biology of the two species. The field body temperature and temperature excess (difference between body and substrate temperature) of two species while being relatively close to each other, showed seasonal differences. Solar radiation, substrate temperature and air temperature were the main effective factors on thermal biology in the field. Additionally, although body size did not have a direct significant effect on body temperature or temperature excess, the interaction between body size and wind were effective on temperature excess. In conclusion, our study partially supports the conservation of thermal biology of related lizard species.

## 1. Introduction

Temperature is one of the most important environmental parameters that strongly influence almost all biochemical reactions and thus most physiological processes linked to fitness of organisms (Angilletta, 2009; Hochachka and Somero, 2002; Ortega et al., 2016a; Pörtner, 2002; Somero, 2004). Eventually, body temperature is one of the most effective factors on the performance of ectothermic animals (Gaitán-Espitia et al., 2014; Huey and Kingsolver, 1989; Huey and Pianka, 2018; Huey and Stevenson, 1979). Thermal performance of ectotherms generally conforms to an asymmetric reaction-norm curve where optimal performance peaks between maximal and minimal thresholds (Angilletta, 2006; Chown et al., 2010; Huey and Stevenson, 1979; Kingsolver and Huey, 1998; Rezende and Bozinovic, 2019; Schulte et al., 2011). Organisms strive to keep their body temperature close to this optimum by thermoregulation to maximize performance in life history including growth, reproduction, foraging, locomotion (Angilletta, 2009; Aparicio Ramirez et al., 2020; Lailvaux and Irschick, 2007; Rutschmann et al., 2020; Seebacher and Franklin, 2005).

Body temperatures of ectotherms, including reptiles, strongly depend on thermal properties of their microhabitats (Goller et al., 2014; Gómez Alés et al., 2017; Huey and Slatkin, 1976; Ortega and Pérez-Mellado, 2016). However, terrestrial environments show thermal variation to varying degrees in several spatial and temporal levels, including altitudinal (Kuyucu et al., 2018; Lymburner and Blouin-Demers, 2020; Van Damme et al., 1989; Zamora-Camacho et al., 2016), seasonal (Díaz et al., 2006; Huey and Pianka, 1977; Liz et al., 2019; Van Damme et al., 1987) and microhabitat level (Adolph, 1990; Goller et al., 2014; Ortega et al., 2016b; Scheers and Van Damme, 2002). Because of this heterogeneity of the thermal environment, behavioral thermoregulation holds crucial importance for ectothermic organisms including reptiles (Cowles and Bogert, 1944; Huey and Slatkin, 1976; Kearney et al., 2009; McMaster and Downs, 2013; Rangel-Patiño et al., 2020). The thermoregulatory strategies vary both among and between species, which depends on the costs and benefits of thermoregulation which also depend on several biotic and abiotic factors in the environment (Huey and Slatkin 1976, Lymburner and Blouin Demers 2020). Species that rely on behavioral thermoregulation can utilize solar radiation by basking (heliothermy) or regulate their heat exchange with the substrate (thigmothermy) (Belliure and Carrascal, 2002; Garrick, 2008; Penniket and Cree, 2015; Rutschmann et al., 2020) to keep their body temperature in set point limits. Most Lacertids are classified as heliothermic, since they effectively shuttle between sun and shade in order to keep their body temperature (*T_b_*) in certain optimum intervals (Basson et al., 2017; Díaz et al., 1996; Van Damme et al., 1987).

It is shown that even sympatric squamate species that live in the same environments might differ in their usage of thermal micro-habitats (Angilletta, 2009; Gomes et al., 2016; Gómez Alés et al., 2017). Therefore, differentiations in thermoregulatory strategies can also result in alterations for their habitat use in spatial and temporal scales (Buckley et al., 2015; Lymburner and Blouin-Demers, 2020; Neel and McBrayer, 2018; Paterson and Blouin-Demers, 2018). Microhabitat selection (Castilla and Bauwens, 1991; Ortega and Pérez-Mellado, 2016), reproduction activities (Ibargüengoytía et al., 2008; Luo et al., 2010; Meiri et al., 2012; Wang et al., 2016), foraging rates (Kearney et al., 2020; Verwaijen and Van Damme, 2007) and growth patterns (Angilletta et al., 2002; Have and Jong, 1996; Refsnider et al., 2019; Sears et al., 2016) are partly determined by thermal factors in ectotherms’ environment. The heterogeneity of the spatial thermal environment has an important role in thermoregulation (Logan et al., 2019, 2015; Sears et al., 2011). This heterogeneity provides more opportunities for activities and regulations for lizards by enabling lizards to behaviorally regulate their energetic thermoregulation costs (Ortega and Martín-Vallejo, 2019).

Air temperature, substrate temperature, humidity, solar radiation, wind speed, availability of shade can be assessed as some of the most important environmental factors that influence thermal activities of lizards (Fei et al., 2012b; Kiefer et al., 2007; Logan et al., 2015). In addition to abiotic factors in the environment morphological and physical characteristics can also influence the thermal biology of ectotherms, such as coloration and body size (Clusella-Trullas and Chown, 2014; Sagonas et al., 2013a; Tanaka, 2005). For instance, darker or melanic coloration might increase the amount of solar radiation absorbed by integument, providing potential advantage to darker colored ectotherms in colder environments (Castella et al., 2013; Clusella-Trullas et al., 2008; Matthews et al., 2016; Stuart-fox et al., 2017). Another important factor on heat exchange is body size, case studies on ectotherm species have shown that generally there is a negative correlation between body size and heat exchange rate (Belliure and Carrascal, 2002; Herczeg et al., 2007; Lactin and Johnson, 1997; Rutschmann et al., 2020), i.e larger individuals heat up more slowly compared to smaller ones by absorbing solar radiation and conduction with hot substrates (Belliure and Carrascal 2002), however this might also lead to losing heat in a slower rate in colder environments (Garrick, 2008). Small size also can be both an advantage and disadvantage, for example it might enable smaller ectotherms to heat up more rapidly to optimal temperatures, which is particularly useful for lizards under predator pressure and should move quickly when predation danger is present, on the other hand this higher heating rate carries with it the danger of overheating and individuals have to retreat quickly to shelters before body temperatures reaches harmful levels under very high temperatures (Gardner et al., 2011). Additionally, individuals with larger body size are able to maintain their body temperature more efficiently in shelter after basking, needing less shuttling between different thermal spaces, reducing the exposure time to predators (Dzialowski and O’Connor, 2001; O’Connor et al., 2000; Sagonas et al., 2013a), on the other hand large ectotherms are also sensitive to overheating in hot and open environments as they also cool down slower (Maia-Carneiro and Rocha, 2013; Porter and Gates, 1969). Wind speed is another important parameter that might be influential on body temperature of ectotherms due to heat loss by convection, and high wind speed might significantly reduce the effectiveness of thermoregulation (Maia-Carnero et al. 2012, Logan et al. 2015, Ortega et al. 2017).

Here, we study and compare the thermal biology of two sympatric lizards in western Anatolia with emphasis on important biotic and abiotic variables in the environment, including body size, sex, air temperature, substrate temperature, solar radiation and wind speed. We monitored *Lacerta diplochondrodes* and *Parvilacerta parva* in Phrygian valley during their active season by several *in situ* observations. Thermoregulation dynamics of various lacertid species were studied in mainland and islands of Greece in the last decade (Sagonas et al. 2013 a,b). However, there is a serious lack of field-based thermal ecology studies on not only our model species but also for all terrestrial reptiles in the Anatolian Peninsula. Therefore, three questions were set for this work: i) Do these two species differ in their thermal biology? ii) To what degree body size is effective on this potential difference. iii) Which microclimatic variables are effective on the thermal biology of these two species and to what extent?

## 2. Materials and methods

### 2.1 Model organisms

*Lacerta diplochondrodes* Wettstein, 1952 (formerly known as *Lacerta trilineata* in Western Anatolia) a large-sized lizard of Lacertidae, is evaluated to be endemic to the Anatolian Peninsula after recent phylogeographic queries (Kornilios et al., 2020). This large-sized Lacertid can reach up to 50 cm as a body length. The dorsal colour is brownish and there are 5 light coloured longitudinal lines present in juveniles. There may also be a dashed line on both sides of the ventralia. As the lizard grows the color of dorsalia turns to green, light lines disappear and small and dark spots occur densely. These spots are generally more common in females. Light blue coloring is observed on both sides of the head in males. The ventralia is yellowish white in males and pinkish yellow in females. It lives in steppe habitat, where shrubs and rocks are abundant. Despite its large size, its movements are very fast and it tends to hide between the roots of shrubs.

The smaller sized *Parvilacerta parva* (Boulenger, 1887) inhabits steppe vegetation along inner parts of Anatolian Peninsula and Armenian mountainous steppes. It is a dwarf lizard with a body length of up to 14 cm. The dorsal plates are keeled, and the coloration of this part is grayish or light brown with black and white spots. Similarly, there are also spots on lateral sides of the body. While the coloration of ventralia is yellowish or white in males, it is generally white in females. Therefore, the highlighted dimorphic morphological character is the presence of blue spots on ventral laterals in only males.

### 2.2 Study area

Although the entire Phrygian region covers parts of Ankara, Afyonkarahisar and Eskişehir provinces, plus most of Kütahya province and northern slopes of Konya, Isparta and Burdur in Western Anatolia, the study area range is only northern slopes of this region in Afyonkarahisar (Lat: 39.046638, Lon: 30.522929, Elevation: 1145 m) (Fig. 1). There are many deep valleys in this slope and elevation varies between 1100 and 1500 meters. The rock formations are shaped from soft volcanic tuffs, which are dated to Tertiary Era (Gürsoy et al., 2003). This also allows metamorphism of the greywackes and their sedimentation by corrosion process causes alluvial soil type. The microclimate provided by this soil type has its own characteristics: Hot and dry in summers and cool and less humid in winters. Vegetation in the Phrygian Valley is highly dominated by floristic steppe elements. The study area, which is the northern slopes of valley, includes various annual *Poaceae* species, that covers almost the entire area. Apart from that, shrubs are abundant in several areas, where *L. diplochondrodes* specimens were observed. However, due to Valley’s geological status tuff based bedrock landscape is surrounding the study area (Gürsoy et al., 2003).

**Figure 1.**
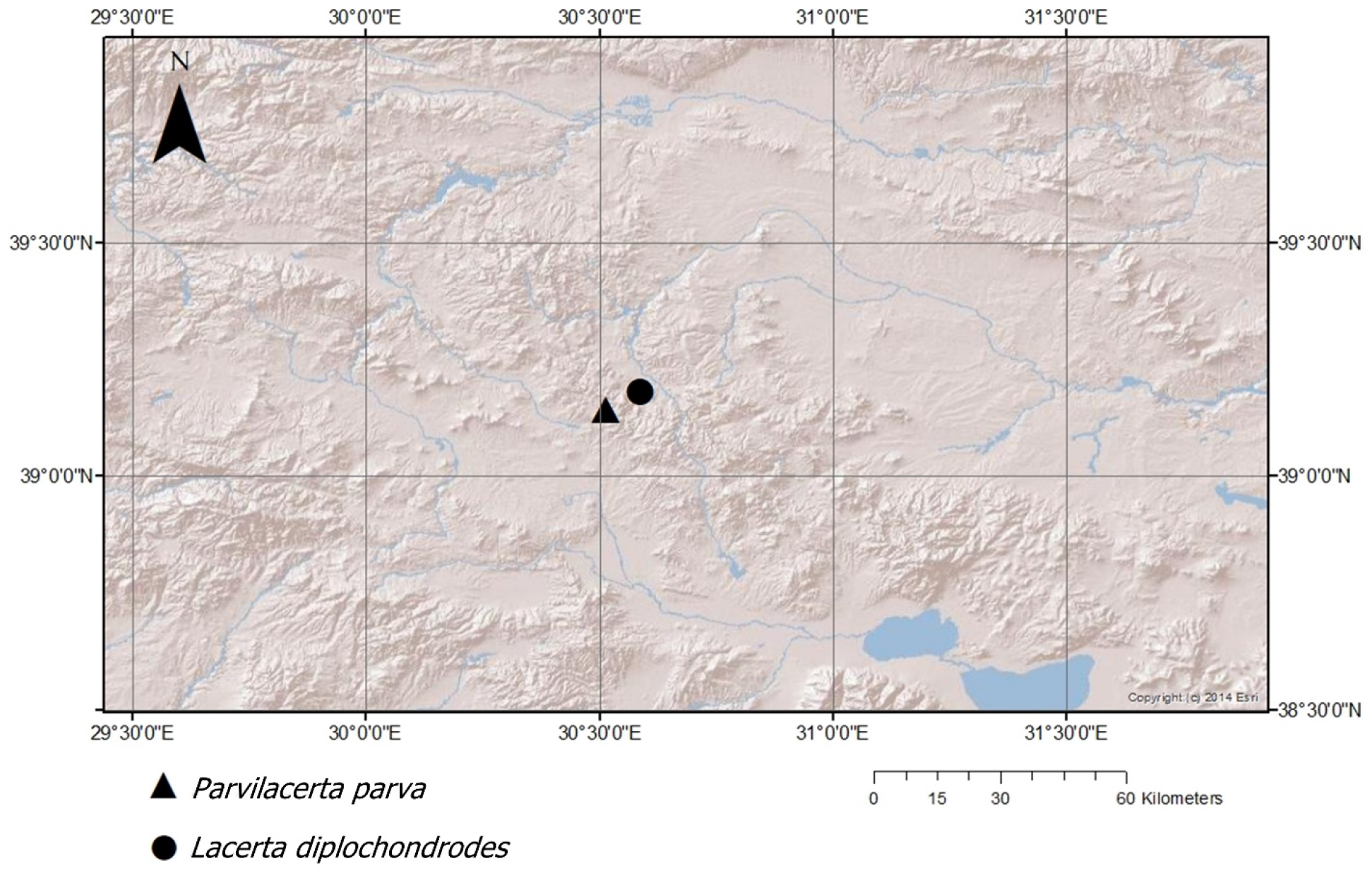
Study area: Phrygian Valley, Afyon, Turkey

The mean annual temperature in the study area is 10.2° C and as a result of terrestrial climate, the hottest months are July (22.1° C) and August (20.6° C) (TSMS, 2020). The precipitation is high in spring but mean precipitation is still low due to two factors: i) distance from the sea side, ii) low humidity. The mean annual precipitation is 406.1 mm and the cloudiness rate is 4.7/10 (TSMS, 2020). Therefore, according to Köppen-Geiger climate classification, the climatological conditions of the study area displays “Csa” pattern (Warm climate with hot dry summer) (Beck et al., 2018; Turkes, 2020).

### 2.3. Field Sampling

Lizards from both species were simultaneously studied between May and September 2018 for 45 non-rainy days (9 days in May, 7 days in June, 5 days in July, 15 days in August and 9 days in September). In order to evaluate the activity and variation in body temperature of lizards, they were captured from 07.00 to 18.00 during their diurnal activity times by visual encounter survey (VES) method (Maia-Carneiro et al., 2012), an effective method for capturing the lizards which live in open habitats and are locally abundant.

The body temperatures (*T_b_*) were measured with an infrared thermometer (Fluke 62 MAX). Due to their low invasiveness, non-contact methods such as infrared thermometers and infrared thermography are now widely used in thermal biology studies on reptiles (Barroso et al., 2016; Carretero, 2012; Tattersall, 2016). The measurements were taken from lizards’ dorsal part between their head and neck immediately 2-5 seconds after capture to avoid heat transfer to animal (0.1°C sensitivity). The hemipenis presence/absence is also an easy way for the sexual identification of specimens. Total body length (BL) was measured from snout to the tip of the tail by a digital caliper (Ecotone ^®^, to the nearest 0.1 mm). After these examinations, the lizards were released to the same point where they were originally captured.

Air temperatures (*T_a_*) were taken with a digital thermometer (HTC 288-ATH ^®^). Substrate temperatures (*T_s_*) were taken with the same digital infrared laser thermometer (Fluke Model 62 ^®^) that was used for body temperature measurements from the lizard capture point. Substrate type was recorded from the lizard capture point as grass, rock or soil. Wind speed (m/s) was measured with a Windtronic 2 ^®^ anemometer and solar radiation (sol) (w/m^2^) was measured with an Apogee ^®^ pyranometer.

All field works were done under permission of the Ministry of Agriculture and Forest (27.02.2018, 72784983-488.04-49994) and Hacettepe University Local Ethical Committee (31.01.2018/52338575-4).

### 2.4. Statistical Analysis

We built two ordinary least square linear models for *T_b_* and *T_ex_*. Compliance to normal distribution of parameters were checked with Shapiro-Wilks test. The distribution of *T_b_* did not conform to a normal distribution, approaching a more leptokurtic distribution so we used a bijective inverse transformation using the lambertW package for R (Goerg, 2020) to transform the data to a normal distribution. *T_b_* was taken as the dependent variable and species, sampling month (May to September), sampled substrate (soil, grass and rock) and sex (juvenile, male and female) were determined as categorical predictors while body size (measured as body length in mm), air temperature (*T_a_*), substrate temperature (*T_s_*), solar radiation (w/m^2^) and wind speed (m/s) were continuous predictors. Additionally, we included the following interactions, body length (BL): solar radiation, body size: wind speed, species: month and species: substrate. As the distribution of *T_ex_* conformed to a normal distribution, we built the linear model without data transformation to compare the effects of same environmental and morphological parameters on temperature excess including interactions, except *T_a_* which was left out which since it was directly related with *T_ex_*. After building the initial models we used AIC (Akaike Information Criteria) (Aho et al., 2014; Burnham and Anderson, 2004; Symonds and Moussalli, 2011) backward stepwise elimination to gather the most informative model. Variance inflation was also checked and variables with VIF > 4 were excluded from the final models to minimize collinearity. The substrate use between species were compared by building cross tables for proportion of observations in three substrate types and comparisons were made by chi-square test. All statistical analyses and transformations were carried out in R statistical software (R. Core Team 2018).

## 3. Results

On general both mean *T_b_* (33.7 ± 0.13 °C) and *T_ex_* (9.38 ± 0.14 °C) of the smaller (BL = 114.59 ± 0.84 mm) *P. parva* was higher compared to larger *L. diplochondrodes* (*T_b_*: 32±0.12 °C, *T_ex_*: 8.99 ± 0.18 °C, BL = 326.2 ± 2.55 mm) (Table 1). As expected, body temperature reached its highest points during the middle of summer for both species. It can be seen from Table 2 and Fig 2 that temperature excess parameters show a monthly fluctuating pattern, as for both species *T_ex_* decreased between May and June, increasing again between June and July and decreasing after, on the other hand although *T_ex_* continued to decrease for *Parvilacerta*, for *Lacerta diplochondrodes*, it increased again in September (Table 2). According to the final model for body temperature, sampling month (September), type of substrate, *T_a_, T_s_* and solar radiation were significant factors. Additionally, although species was not a significant factor the interaction between species (*P. parva*) and substrate (rock) had significant effect (Table 3). For the temperature excess model, significant factors were sampling month, substrate (rock) and solar radiation. Additionally, we found that interactions species: month, species: substrate and body size: wind speed had significant effect on temperature excess (Table 4). The interaction between species and months i.e. the differential pattern of temperature excess is apparent in Fig. 3. We also detected that the two species differed in habitat use where *P. parva* had higher preference for grassy habitat (54% grass, 42% soil, 3% rock) whereas *L. diplochondrodes* had higher preference for open soil (29% grass, 69% soil, 1% rock) (χ^2^ = 77.25, df = 2, *p* < 0.001).

**Figure 2.**
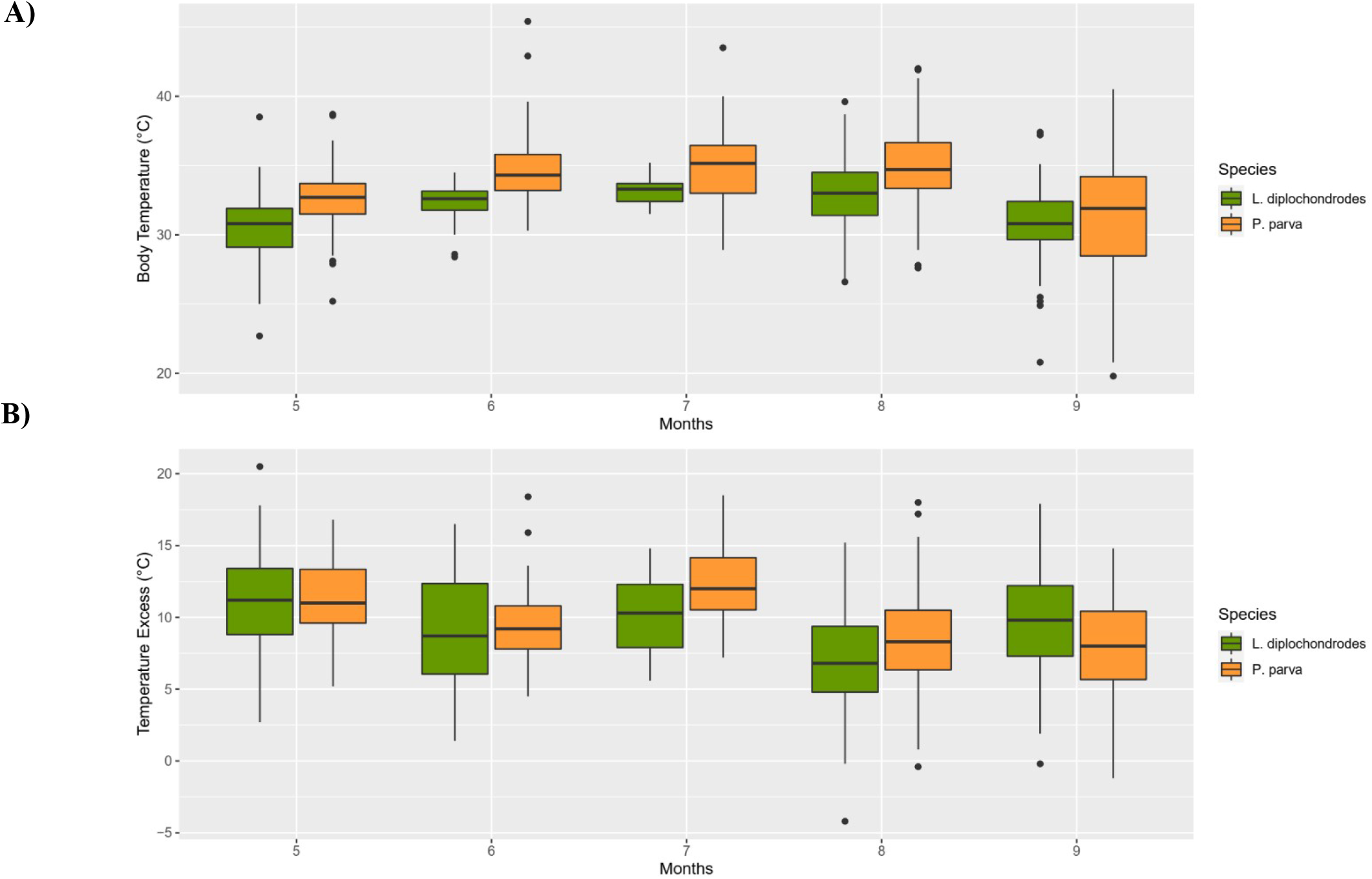
**A)** Monthly boxplots of body temperature (*T_b_*) values for *L. diplochondrodes* and *P. parva*. **B)** Monthly boxplots of temperature excess (*T_ex_*) values for *L. diplochondrodes* and *P. parva*.

**Figure 3.**
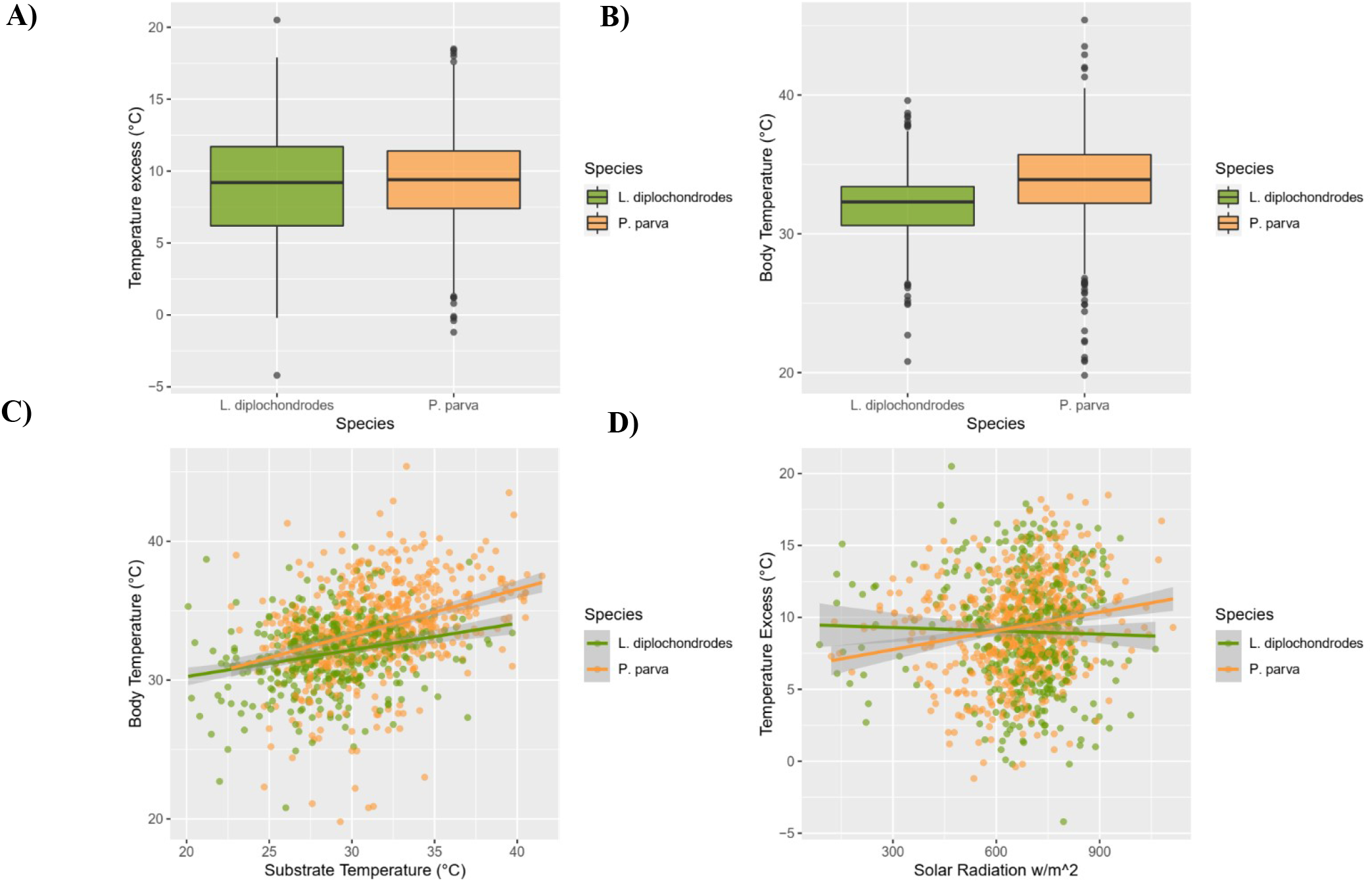
**A)** Boxplots of temperature excess (*T_ex_*) values for *L. diplochondrodes* and *P. parva*. **B)** Boxplot values of body temperature (*T_b_*) values for *L. diplochondrodes* and *P. parva*. **C)** Scatteplot of body temperature (*T_b_*) with substrate temperature (*T_s_*). **D)** Scatteplot of temperature excess (*T_ex_*) with solar radiation (w/m^2^).

**Table 1.** Mean body temperature (T_b_), mean temperature excess (T_ex_) and mean body length of two lizard species.

| Species | Mean $T_b \pm$ St. error | $T_{ex} \pm$ St. error | BL ± St. error |
| --- | --- | --- | --- |
| *Lacerta diplochondrodes* | $32 \pm 0.12$ °C | $8.99 \pm 0.18$ °C | $326.2 \pm 2.55$ mm |
| *Parvilacerta parva* | $33.7 \pm 0.13$ °C | $9.38 \pm 0.13$ °C | $114.59 \pm 0.84$ mm |

**Table 2.** Monthly Mean body temperature (T_b_) and mean temperature excess (T_ex_) of two lizard species.

| Species | Month | $T_b \pm$ St. error | $T_{ex} \pm$ St. error | N |
| --- | --- | --- | --- | --- |
| *Lacerta diplochondrodes* | May | $30.6 \pm 0.25$ °C | $11.1 \pm 0.37$ °C | 91 |
| | June | $32.4 \pm 0.14$ °C | $9.1 \pm 0.47$ °C | 70 |
| | July | $33.1 \pm 0.13$ °C | $10.4 \pm 0.36$ °C | 49 |
| | August | $32.9 \pm 0.21$ °C | $6.83 \pm 0.27$ °C | 150 |
| | September | $30.9 \pm 0.28$ °C | $9.62 \pm 0.41$ °C | 90 |
| *Parvilacerta parva* | May | $32.6 \pm 0.19$ °C | $11.3 \pm 0.24$ °C | 119 |
| | June | $34.7 \pm 0.25$ °C | $9.35 \pm 0.25$ °C | 89 |
| | July | $34.9 \pm 0.34$ °C | $12.4 \pm 0.34$ °C | 58 |
| | August | $34.9 \pm 0.18$ °C | $8.34 \pm 0.22$ °C | 223 |
| | September | $31.4 \pm 0.39$ °C | $7.97 \pm 0.3$ °C | 120 |

**Table 3.**
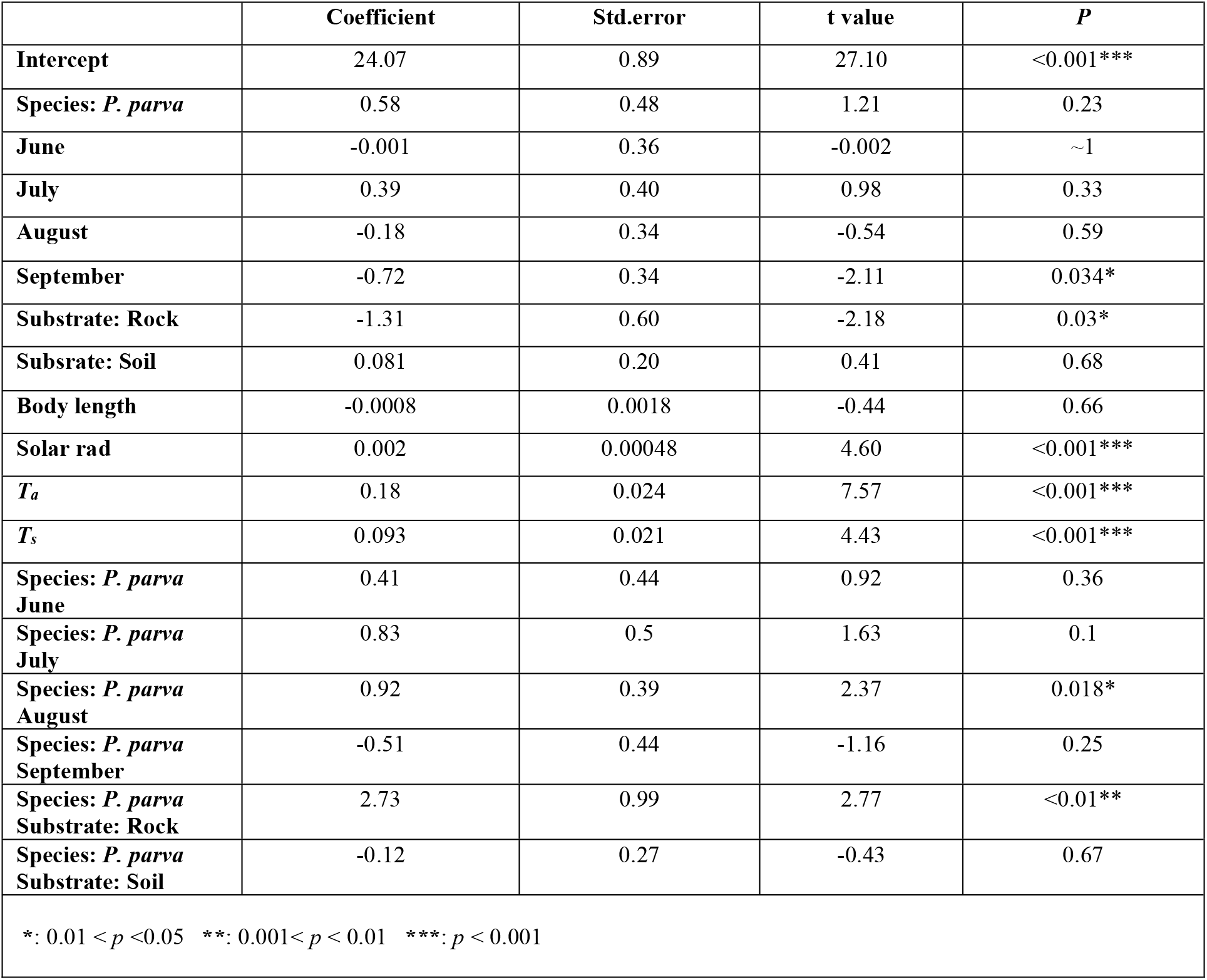
Results of the best model for parameters effective on body temperature (*T_b_*).

**Table 4.**
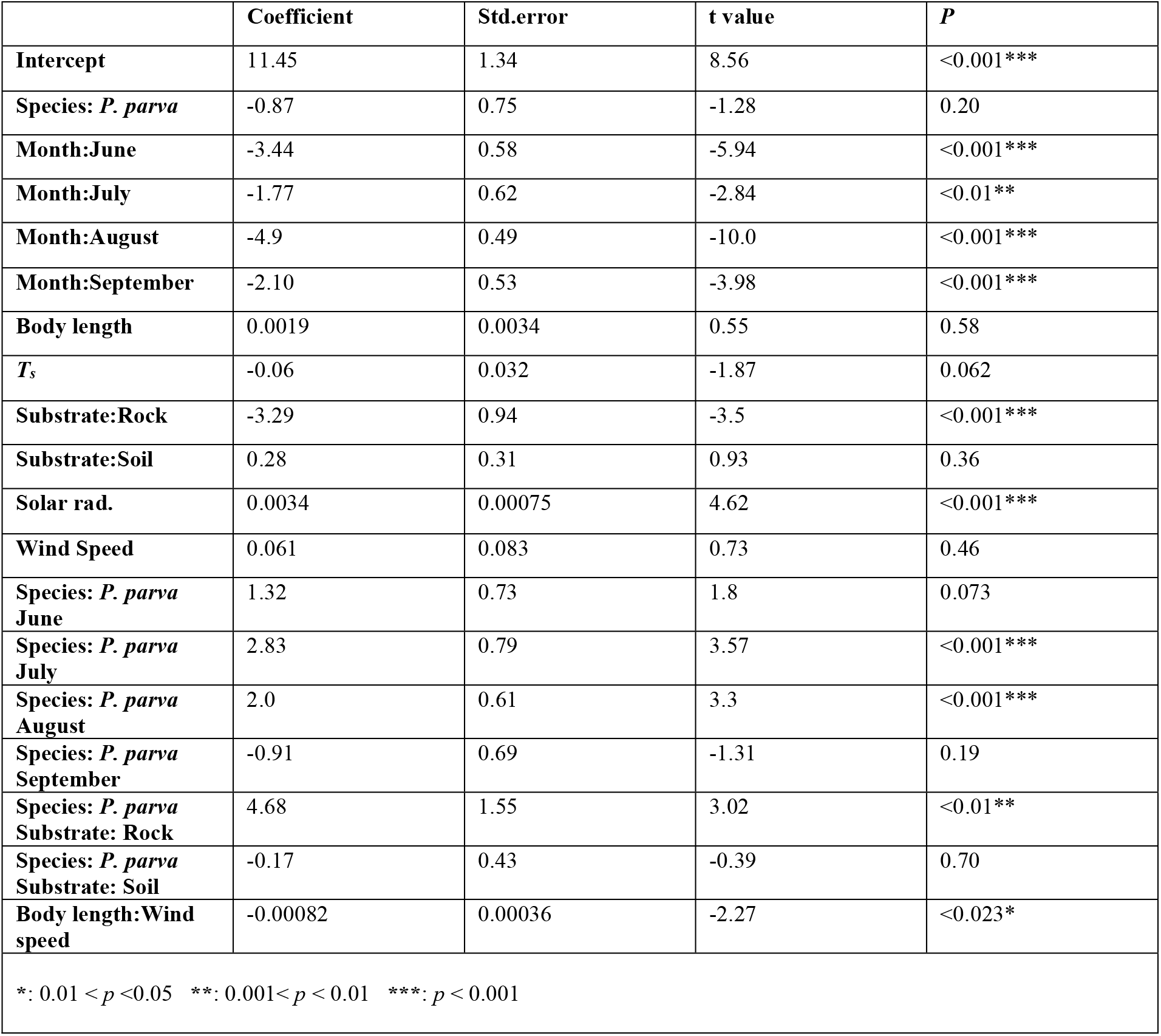
Results of the best model for parameters effective on body temperature (*T_ex_*).

|  | Coefficient | Std.error | t value | P |
| --- | --- | --- | --- | --- |
| **Intercept** | 11.45 | 1.34 | 8.56 | <0.001*** |
| **Species: *P. parva*** | -0.87 | 0.75 | -1.28 | 0.20 |
| **Month:June** | -3.44 | 0.58 | -5.94 | <0.001*** |
| **Month:July** | -1.77 | 0.62 | -2.84 | <0.01** |
| **Month:August** | -4.9 | 0.49 | -10.0 | <0.001*** |
| **Month:September** | -2.10 | 0.53 | -3.98 | <0.001*** |
| **Body length** | 0.0019 | 0.0034 | 0.55 | 0.58 |
| **$T_s$** | -0.06 | 0.032 | -1.87 | 0.062 |
| **Substrate:Rock** | -3.29 | 0.94 | -3.5 | <0.001*** |
| **Substrate:Soil** | 0.28 | 0.31 | 0.93 | 0.36 |
| **Solar rad.** | 0.0034 | 0.00075 | 4.62 | <0.001*** |
| **Wind Speed** | 0.061 | 0.083 | 0.73 | 0.46 |
| **Species: *P. parva***<br>**June** | 1.32 | 0.73 | 1.8 | 0.073 |
| **Species: *P. parva***<br>**July** | 2.83 | 0.79 | 3.57 | <0.001*** |
| **Species: *P. parva***<br>**August** | 2.0 | 0.61 | 3.3 | <0.001*** |
| **Species: *P. parva***<br>**September** | -0.91 | 0.69 | -1.31 | 0.19 |
| **Species: *P. parva***<br>**Substrate: Rock** | 4.68 | 1.55 | 3.02 | <0.01** |
| **Species: *P. parva***<br>**Substrate: Soil** | -0.17 | 0.43 | -0.39 | 0.70 |
| **Body length:Wind**<br>**speed** | -0.00082 | 0.00036 | -2.27 | <0.023* |
| *: 0.01 < $p$ < 0.05 **: 0.001 < $p$ < 0.01 ***: $p$ < 0.001 | | | | |

## 4. Discussion

Here we studied the thermal biology of two sympatric Lacertid species (*L. diplochondrodes* and *P. parva*) in Phrygian Valley, Turkey. As expected for heliothermic lizards that depend on basking for thermoregulation; air temperature (*T_a_*), substrate temperature (*T_s_*), seasonality (months) and solar radiation were the most effective parameters on both body temperature and temperature excess (Clusella-Trullas et al., 2008; Garrick, 2008; Matthews et al., 2016). On the other hand, body size was not a statistically significant factor in *T_b_* and *T_ex_* by itself. Although numerous studies (especially those which successfully implement detailed biophysical models into experimental study frames) documented the significant effect of body size on heating and cooling rate of lizards (Fei et al., 2012a; Garrick, 2008; Herczeg et al., 2007; Shine and Kearney, 2001), this relation might not be detected directly in field conditions where several other important environmental, morphological and behavioral factors can conceal this interaction (Christian et al., 2006; Corkery et al., 2018; Sagonas et al., 2013a; Stevenson, 1985). First, the effect of body size is a two way process as increased thermal inertia slows both the heat gain and heat loss from ectotherm bodies (Garrick, 2008; Herczeg et al., 2007; Maia-Carneiro and Rocha, 2013; Zamora-Camacho et al., 2014). On the other hand, despite not observing a substantial relation between either body size or wind speed and *T_ex_*, we detected a significant negative effect of the interaction between body size and wind speed on *T_ex_*. This finding points that heat loss by convection of a larger area of exposed body surface might be an important factor on thermal biology in the field. Previously, it was shown that increased wind speed can significantly decrease the effectiveness of thermoregulation and impend activity in exposed open high elevation habitats (Huang et al., 2020; Logan et al., 2015; Maia-Carneiro et al., 2012; Ortega et al., 2017). Our study area is an open steppe that is relatively more exposed to wind effects with low bushes and grass as vegetation, additionally the wind speed in the area was relatively high during the measurements (mean wind speed: 2-5 m/s). Added together these components might be increasing the effect of wind speed on body temperatures in the field.

Although observation of gender differences in thermal biology of reptiles are relatively rare, gender as a factor should be taken into consideration due to the biological importance of these rare cases (Huey and Pianka, 2007). A proportion of cases where significant difference observed for thermal biology this variation can be connected with sexual size dimorphism (Lailvaux, 2007; Ortega et al., 2016b). On the other hand thermal biology differences between genders might also be connected with other gender specific differences in performance and behavior such as locomotion, stamina, assimilation efficiency and micro-habitat preference (Beal et al., 2014; Lailvaux et al., 2003). In our study there was not any visible sexual size dimorphism for both species and we did not detect any significant effect of gender or ontogeny (between juveniles and adults) on body temperature or temperature excess. These results suggest that regardless of gender or ontogeny the inviduals belonging to two species might have similar thermoregulatory behaviour in the field.

Although we measured body temperatures from the neck region by infrared thermography our measurements (32 ± 0.12°C) for *L. diplochondrodes* were close to body temperature measurements taken from *L. trilineata*, a very closely related species in mainland Greece [Stymphalia: 31.94±1.65°C, Karditsa: 32.30±1.54°C st.dev, (Sagonas et al., 2013c)]. Additionally, the relatively close pattern of body temperatures across different months suggest that these two species maintain similar body temperatures to each other through variable environmental temperatures. These results are in compliance with results from other studies and reviews, which suggest that thermal biology of lizards is a relatively conserved trait across different species (Clusella-Trullas and Chown, 2014; Gómez Alés et al., 2017; Grigg et al., 2013; Rowe et al., 2020).

It was shown that sympatrically coexisting lizards might differentiate their niches, and this also might affect their thermoregulatory behavior (Akashi et al., 2016; Gomes et al., 2016; Gómez Alés et al., 2017; Sagonas et al., 2017). In our study study we have found that the two species differed in their preferences for different habitat types. On the other hand, we did not detect a significant relation between thermal biology and preference of grass and soil substrates, which are the main habitat types in the study area. The smaller *P. parva* might be preferring grassy habitats for the suitability to hide smaller lizards from predators. The main difference for *T_b_* and *T_ex_* between species were seasonal as shown by the effect of the interaction between month and species, thus our results confirm that seasonality is the main factor on the field body temperature of lacertids, considering that other abiotic parameters including temperature and solar radiation are also highly dependent on seasonality (Ortega and Pérez-Mellado, 2016). Additionally, aside from the direct effects of seasonality on body temperature other important life history activities such as reproduction, foraging and feeding depend on both thermal biology and seasonality patterns (Adolph and Porter, 1996; Christian et al., 2003; Van Damme et al., 1987). This seasonal difference was much more pronounced for *T_ex_* as *P. parva* shows higher *T_ex_* for both July and August with high significance (Table 1) while for *T_b_* this effect was marginally significant for August (Table 2). The relatively stable state of *T_b_* compared to the difference in *T_ex_* between months indicates that the role of behaviour in maintenance of optimal body temperature might be important, as the higher amount of difference between *T_b_* and *T_a_* in warmer periods of year might indicate a higher frequency of overheating avoidance (Vicente Liz et al., 2019). Given that both of these two species are diurnal with most active times being spring and summer, efficient thermoregulatory behaviour is crucial for heat avoidance while maintaining an optimal body temperature for foraging and metabolism (Huey and Pianka, 2018; Sepúlveda et al., 2014). It is also striking that while smaller *P. parva* has higher T_ex_ than *L. diplochondrodes* in July when ground temperature is at its highest, this picture is reversed in September when temperatures are lower. Since we did not detect a direct effect of body size on temperature excess this pattern might have been caused by the temporal behavioral differences between the two species.

Regarding the usage of substrate in the study area, the effect of rock type of substrate was significant and negative on *T_b_* and *T_ex_*. However, the interaction between *P. parva* and rock substrate was significant and positive. Which might be due to the effect of rocky substrate being smaller on *P. parva*, on the other hand both of these species rarely prefer rock, as we observed the individuals from two species on rocky substrates only in 22 times out of 1059 total observations. In addition rocky substrates in the study area were relatively rare with most of the area being either shrubs or naked soil.

Although thermal ecology of various lizard species was investigated in Aegean Islands and mainland Greece in the east Mediterranean region (Sagonas et al., 2017, 2013c, 2013b) to our knowledge this study is the first to investigate the thermal biology of lizard species occurring in Anatolia. On the other hand, additional studies on the operative temperatures (Hertz et al., 1993; Ortega et al., 2014; Sagonas et al., 2013a), voluntary and critical thermal limits (Aparicio Ramirez et al., 2020; Camacho and Rusch, 2017; Qu et al., 2011) and also thermal preferences (Goller et al., 2014; Van Berkel and Clusella-Trullas, 2018; Wang et al., 2013) are necessary for a more realized thermal ecology framework for these species. We believe that this study will also be a first step in constructing a framework for predicting the responses of reptile populations inhabiting western Anatolia to climate change. As thermal traits, particularly thermal behavior and tolerance plays a significant role in determining the responses of ectothermic species to global environmental change (Buckley et al., 2015; Enriquez-Urzelai et al., 2020; Kearney et al., 2009), knowledge of the thermal ecology and physiology differences and microhabitat preferences would provide useful information to infer and prevent the effects of climate change on reptile species under threat (Ceia-Hasse et al., 2014; Kearney et al., 2009).

## Acknowledgments

M.K. Sahin contributed to the study design, collected the field data and contributed to the manuscript draft. A.C. Kuyucu designed and supervised the study, carried out the statistical analysis and drafted the manuscript. This work was supported by Hacettepe University Scientific Research Coordination Unit. (Project Number FHD -2018-16903).

## Data accessibility

Data supporting this study are provided in the link, doi:10.17632/g868g5m438.1

